# Robust constrained weighted least squares for in vivo human cardiac diffusion kurtosis imaging

**DOI:** 10.1101/2025.02.12.637822

**Authors:** Sam Coveney, Maryam Afzali, Lars Mueller, Irvin Teh, Filip Szczepankiewicz, Derek K. Jones, Jurgen E Schneider

## Abstract

**Summary:** Robust estimation with convexity constraints significantly improves signal fitting for in vivo human cardiac diffusion kurtosis imaging.

**Purpose:** Cardiac diffusion tensor imaging (cDTI) is an emerging technique to investigate the microstructure of heart tissue. At sufficiently high b-values, additional information on microstructure can be observed, but the data require a representation beyond DTI such as diffusion kurtosis imaging (DKI). cDTI is highly prone to image corruption, which researchers usually attempt to handle with shot-rejection. However, this can be handled more generally with robust estimation techniques. Recent work has also demonstrated the need to perform constrained fitting for DKI, as fitted parameters can otherwise violate necessary constraints on the signal behaviour, causing significant errors in estimated measures.

**Methods:** We developed robust constrained weighted least squares (RCWLS) by combining robust estimation with convexity constraints specifically for DKI. Using in vivo cardiac DKI data from 11 healthy volunteers collected with a Connectom scanner, we tested various combinations of fitting techniques, with/without robustness and with/without constraints.

**Results:** RCWLS was the only tested technique that convincingly showed radial kurtosis to be larger than axial kurtosis for all subjects, something that is expected in myocardium due to increased restrictions to diffusion in the plane perpendicular to the primary myocyte direction. RCWLS also showed the best correction of corrupted regions in diffusion parameter maps for individual subjects.

**Conclusion:** Fitting techniques utilizing both robust estimation and constraints are essential to facilitate applicability of in vivo cardiac DKI.

## 1 Introduction

Cardiac diffusion weighted imaging (cDWI) is a magnetic resonance imaging (MRI) technique that can be used to investigate cardiac tissue microstructure. Cardiac diffusion tensor imaging (cDTI) is the most common cDWI method used on the heart, from which measures such as mean diffusivity (MD) and fractional anisotropy (FA) can be derived. However, the diffusion-weighted signal in tissue deviates from monoexponential decay at higher diffusion weighting (as expressed by the b-values) due to cell membranes and other restrictions in biological tissue [1–3]. Diffusion kurtosis imaging (DKI) can quantify these deviations. Non-Gaussian diffusion models (including DKI) have been shown to have a higher sensitivity for the detection of hypertrophy in ex vivo rat hearts compared with DTI [3]. Kurtosis measures include mean kurtosis (average kurtosis across all directions), axial kurtosis (kurtosis in the primary diffusion direction) and radial kurtosis (average kurtosis in the plane perpendicular to axial kurtosis) [4–12]. Anisotropic and isotropic kurtosis can be also distinguished with q-space trajectory imaging [13].

Spin echo-based DKI in the human heart in vivo is challenging due to low SNR, a short myocardial T2 (approximately 46 ms at 3.0 T [14]) and long echo-times (TE), required to achieve sufficient motion compensation and b-values. Nonetheless, acquiring data has been shown to be feasible in healthy volunteers in vivo using ultra-strong gradients (i.e. 300 mT/m) at echo times and resolutions comparable to those commonly used for conventional cDTI [15, 16]. However, even for the brain, which has longer T2 and significantly less motion, DKI is challenging: fitting methods need to handle data corruptions [17], but also need to yield a physically plausible signal if kurtosis measures are to be meaningful [18].

Image corruption is a common problem in cDWI [19]. Also, motion causes additional signal variations sometimes referred to as physiological noise. In cDTI, shot-rejection is usually performed in an attempt to handle corruptions where noticeably corrupted images are removed from datasets before fitting, a method that is typically time-consuming and subjective. In our experience, the reduced signal at higher b-values simultaneously causes a larger number of corruptions (including those from misregistration) and a decreased ability to perform shot-rejection effectively. Robust estimation, in which outlier signals are identified and removed at the voxel level, is an alternative to shot-rejection. Our recent work shows that robust estimation is superior to shot-rejection in cDTI [20], so robust estimation in cardiac DKI is worth investigating, but has not been done yet.

Although the DKI signal representation does not correspond to a valid diffusion propagator, the fitted signal should still adhere to the physical principles governing the data-generating process. If it does not, the fitted parameters and any measures derived from them will lack meaningful interpretation. For example, the compartment model predicts that kurtosis should be non-negative [2, 4], and the diffusion tensor should be positive definite as in DTI. For DKI, it is not known how to enforce constraints on kurtosis via reparameterization of the fitting problem, so constraints must consider whether the predicted signal behavior is valid, e.g., in [21], non-linear optimization is (infinitely) penalized if constraints are violated. Recently, linear least squares methods have been developed for enforcing convexity constraints on the cumulant generating function, leading to correction of significant errors in brain DKI [18]. These advantages should also apply to cardiac DKI.

While preventing estimated parameters from violating constraints may be seen as a form of robustness, constrained fitting itself is not inherently robust. This is because the constraints do not change the shape of the fitting cost function in parameter space, which is entirely determined by the data. As a result, outliers can still have a detrimental impact on parameter estimates. Furthermore, although robust fitting may reduce the frequency and impact of constraint violations by removing outlier signals, it cannot guarantee that the parameters that optimize the cost function will not violate the constraints. In this work, we combine robust fitting [20] with convexity constraints [18] using iteratively reweighted least squares (IRLS) to give robust constrained weighted least squares (RCWLS). To our knowledge this is the first time that constrained estimation has been combined with robust estimation in MRI. We apply these methods to in vivo cardiac DKI data collected on a Connectom scanner.

## 2 Methods

### 2.1 Diffusion Kurtosis Imaging

The DKI signal representation can be expressed as [2]:

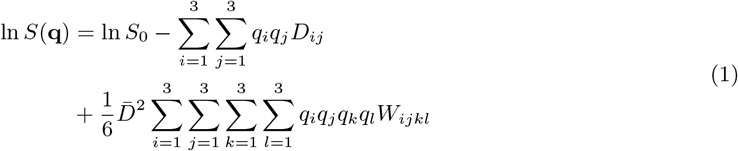

where 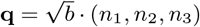 is the (rescaled) wave-vector (denoted this way for convenience in expressing the constraints - see Sec 2.3), and *i, j, k, l* index physical space. The diffusion tensor D and kurtosis tensor W are both symmetric, having 6 and 15 unique elements, respectively. The DTI signal representation is the same as Eq (1) but without the term containing W. Kurtosis is expressed in dimensionless form due to scaling by the mean diffusivity 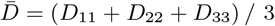.

Expanding Eq (1) accounting for the symmetry of D and W gives the following linear expression:

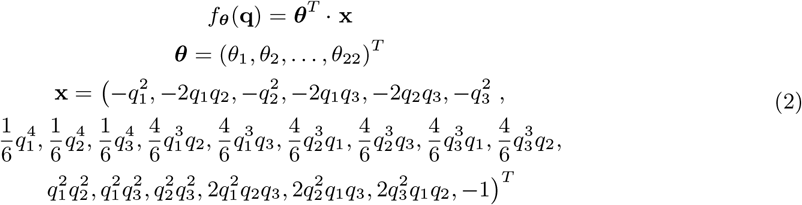

where we have adopted the same sign for the intercept as DiPy [22]. The corresponding expression for DTI would include only terms depending on (*θ*_1_, …, *θ*_6_) and *θ*_22_.

### 2.2 Weighted Least Squares

Given observations *{***q**_**n**_, *S*_*n*_|*n* = 1 … *N}*, the coefficients ***θ*** can be estimated with weighted least squares (WLS):

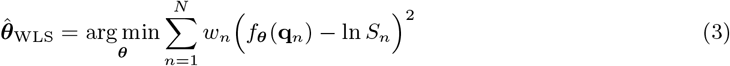

Given design matrix **X** = (**x**_**1**_, **x**_**2**_, …, **x**_**N**_)^*T*^, observation vector **y** = (ln *S*_1_, ln *S*_2_, …, ln *S*_*N*_)^*T*^, and weights vector **w** = (*w*_1_, *w*_2_, …, *w*_*N*_)^*T*^, a weighted design matrix and observation vector can be defined:

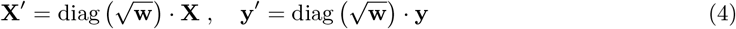

The WLS estimate can then be written as:

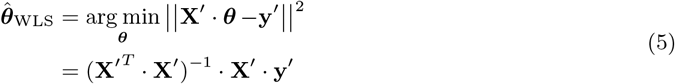

For uniform weights *w*_*n*_ = 1, Eq (5) gives the OLS estimate 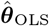. The estimated coefficients can then be related back to the original parameters *D*_*ij*_, *W*_*ijkl*_, and ln *S*_0_.

### 2.3 Convexity Constraints

A useful constraint for DKI is to enforce convexity of the cumulant generating function 𝒞 (**q**) [18]:

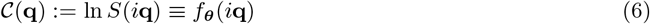

This constraint can be enforced by using sum of squares polynomials [18]; the mathematical background for this can be found in [23]. The semi-definite program for solving the WLS problem subject to constraints, thus yielding the constrained WLS estimate 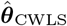, can be written as follows:

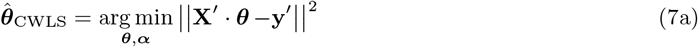

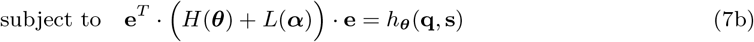

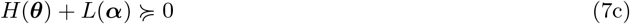

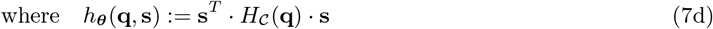

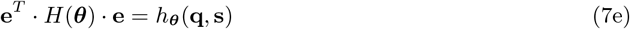

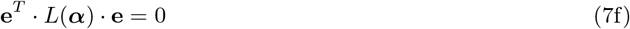

Importantly, we have written the problem in terms of the weighted design matrix **X**^*′*^ and observation vector **y**^*′*^.

We will briefly explain Eq (7) for DKI, leaving details on the constraint matrices *H*(***θ***) and *L*(***α***) (presented here for the first time) for Appendix A. Eq (7d) defines *h*_***θ***_(**q, s**), where *H*_*C*_(**q**) is the Jacobian of *𝒞* (**q**) and the dummy variable **s** has the same dimensions as **q**. Then, *h*_***θ***_(**q, s**) is just a polynomial. Convexity of *𝒞* (**q**) requires that *h*_***θ***_(**q, s**) be non-negative, which can be enforced using a sum of squares polynomial representation, i.e. Eq (7b) (see [23]). We can *exactly* represent *h*_***θ***_(**q, s**) using a relatively small monomial basis for **e** (see Appendix A). The convexity constraint is satisfied when Eq (7c) holds i.e. when *H*(***θ***) + *L*(***α***) is positive semi-definite (PSD). Note that Eq (7) involves optimizing over coefficients ***θ*** and slack parameters ***α***. Appendix B explains how the numerical complexity of the problem can be reduced.

### 2.4 Robust Fitting

In DTI/DKI, WLS usually refers to solving Eq (5) by weighting the (squared) residuals of the linearized problem with the (squared) signal [24]:

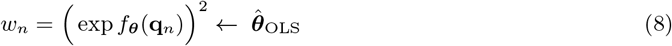

where an initial OLS estimate 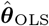 is used to predict the signals (since the true signals are not known). Henceforth, we will use “WLS” to refer to Eq (5) using the weights in Eq (8). Correspondingly, “CWLS” will refer to constrained WLS, i.e. solving Eq (7) using the weights given by Eq (8).

Importantly, neither WLS or CWLS are intrinsically robust, and outlier data can have a detrimental effect on the estimates 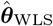 or 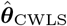. Robust estimation can be implemented using Iteratively Reweighted Least Squeares (IRLS), using weights derived from a robust estimator [17, 20, 25]. Notably, for WLS in DTI/DKI, these robust weight should be chosen so as to preserve the cost function implied by equations (3) and (8) (see [20] for a derivation), such that IRLS solves the WLS/CWLS problem in a robust way. Robust fitting in the DTI/DKI literature is usually done in order to remove the influence of outlier data on the fitted signal, thus making it easier to identify outlier data so that the original problem can be solved without robust weights but also without the outliers.

A robust weighting scheme accounting for the DTI/DKI weights in Eq (8), based on the Geman-McClure M-estimator, with *K* iterations in total, is as follows [20, 25]:

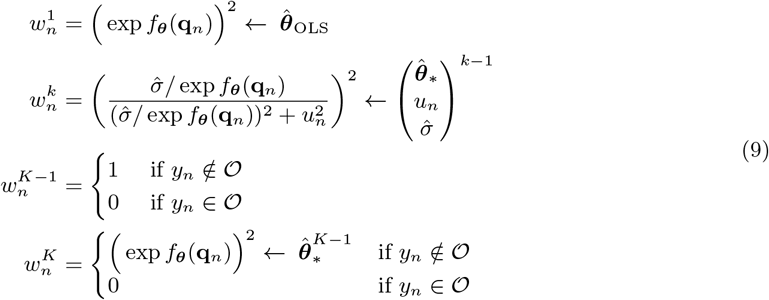

where 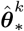 corresponds to the estimated coefficients for iteration *k*. For this scheme, at least 4 iterations would be required. The residuals of the WLS problem are defined as the difference between the log observed signal and log predicted signal:

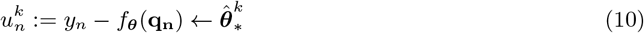

The estimated noise level 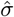 is estimated at the *k*th iteration using a robust estimator designed for the WLS problem [25]:

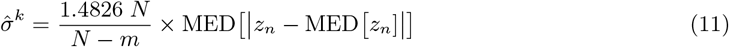

where 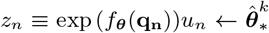, MED is the median operator and *m* is the number of regressors, i.e. *m* = 7 for DTI and *m* = 22 for DKI. The set 𝒪 corresponds to outliers defined by a 3-sigma rule in the signal space:

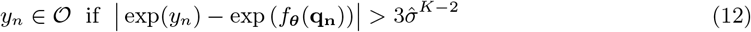

The main insight in this paper is that we can use IRLS with a robust weighting scheme designed specifically for DTI/DKI, i.e., Eq (9), but we are free to choose whether to estimate the coefficients at each iteration using (unconstrained) WLS with Eq (5) or CWLS with Eqs (7). The constraints are independent of the weights, which only enter the cost function Eq (7a) through Eq (4). We will refer to IRLS with weights given by Eq (9) as Robust WLS (RWLS) if Eq (5) is used at each iteration, or as Robust Constrained WLS (RCWLS) if Eq (7) is used at each iteration. The estimated coefficients obtained from the last iteration with weights **w**^*K*^ are denoted as 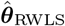 for RWLS and 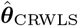 for RCWLS. For convenience, the unconstrained OLS estimate 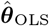 is used to define weights for the first iteration for both RWLS and RCWLS.

We modified DiPy [22] to be able to solve RWLS and RCWLS. These modifications have been incorporated in the latest release (v1.10).

### 2.5 Experimental setup and recruitment

Cardiac diffusion-weighted images (cDWI) were acquired on a Connectom 3T research-only MR imaging system (Siemens Healthcare, Erlangen, Germany) with a maximum gradient strength of 300 mT/m and slew rate of 200 T/m/s. An 18-channel body receive coil was used in combination with a 32-channel spine receive coil. Eleven healthy volunteers (with no known previous cardiac conditions) were recruited for this study: age range 20.5 *±* 1.9 years (18 − 24 years), weight range 64.1 *±* 11.4 kg (54 − 94 kg), 7 females. The study was approved by the local institutional review board (Cardiff University School of Psychology Research Ethics Committee) and all subjects provided written consent.

A prototype pulse sequence was used that enables diffusion encoding with user-defined second-order motion-compensated (M2) diffusion gradient waveforms, designed with the NOW toolbox [26–29]. The maximum gradient strength used in this study for the M2-waveform to generate the b-value of 1350 s*/*mm^2^ was 285.4 mT/m and the maximum physiologically-limited slew-rate was 76.2 T/m/s [15]. The cDWI parameters were: TR = 3 RR-intervals, TE = 61 ms, EPI readout, field-of-view = 320 *×* 120 mm^2^ using ZOnally-magnified Oblique Multislice (ZOOM, tilted RF: excitation, tilt angle: 15^*°*^, tilted slice thickness: 20 mm) [28, 30], in-plane resolution = 2.7 *×* 2.7 mm^2^, slice thickness = 8 mm, 3 short axis slices (base, mid, and apical), partial Fourier factor = 7/8, no parallel imaging, bandwidth = 2354 Hz/pixel. Each full data set was comprised of 5 b-values (b = 100, 450, 900, 1200, 1350 s*/*mm^2^). For *b* ≥ 450 s/mm^2^, 30 directions per shell with 6 repetitions; for *b* = 100 s/mm^2^, 3 directions with 12 repeats. Data were acquired with ECG-gating and under free-breathing. The trigger delay was adjusted for cDWI acquisition in mid-end systole. Saturation bands were placed around the heart. Fat suppression was performed using the SPAIR method [31]. The total acquisition time was approximately one hour.

### 2.6 Post-processing

Post-processing was done using in-house tools [19], with image registration utilizing SITK [32] and fitting utilizing DiPy (with our updates) [22]. Image registration was performed by masking a suitable b = 100 s/mm^2^ image, registering all b = 100 s/mm^2^ images to this reference image, then using the average of registered b = 100 s/mm^2^ images to register the entire dataset. Correlation was used as the registration metric, since it outperformed mutual information for high b-value images. The DTI signal representation was then fit to b ≤ 450 s/mm^2^ images using RWLS, and the full image series was predicted. Each original (unmodified) image was then registered to the corresponding predicted image. This method, similar to [33], improved the registration.

After registration, we fit the DTI signal representation to the b ≤ 450 s/mm^2^ data using RWLS. We calculated MD, FA, and Helix Angle (HA) (using a cylindrical coordinate system with origin on the LV blood-pool center). Segmentation of the LV contours was performed with care taken to exclude voxels exhibiting strong partial-volume effects. For regions strongly affected by artifacts, such as aliasing or susceptibility-induced warping, fitting results do not reliably represent tissue properties. We defined artifact masks using sectors centered on the LV blood-pool, in order to ignore these parts of the myocardium when calculating voxel statistics. We performed this masking by considering the image series, as well as utilizing MD, FA, HA, root-meant-square-error and coefficient of determination (R^2^) from the RWLS DTI fit to *b* ≤ 450 s/mm^2^ data. We have not utilized DKI results to identify artifacts in any way, since this would likely bias comparison between the fitting methods under study.

Having registered the images and segmented the myocardium, we performed further fitting experiments on myocardial voxels only. Defining b_max_ as the maximum b-value images that were utilized in a given fit (such that all images with a lower b-value were also included), we performed the following: DTI using WLS and RWLS for b_max_ = 450 s/mm^2^; DKI with WLS, RWLS, CWLS, and RCWLS, for b_max_ values 900 s/mm^2^, 1200 s/mm^2^, and 1350 s/mm^2^. For RWLS and RCWLS, we used *K* = 10 iterations. We calculated the following measures in each voxel: mean diffusivity (MD), fractional anisotropy (FA), mean kurtosis (MK), axial kurtosis (AK), radial kurtosis (RK), and radial / axial kurtosis (RK/AK) [2,4]. We then calculated the average of these measures over non-artifact myocardial voxels.

## 3 Results

### 3.1 Statistical analysis

Figure 1 shows boxplots of the average DTI/DKI measures for all 11 subjects. All fitting methods are shown for all b_max_ values. The black markers drawn along the bottom of each subplot indicate nonnormality as tested with the Shapiro-Wilk test. Figures 2, 3, and 4 show the results of significance tests between measures from the different fitting methods and different b_max_ shown in Figure 1. The Wilcoxon signed-rank test was used, since non-robust methods were found to often produce results that violate the assumption of normality (see Figure 1). Figure 2 shows the p-values between different methods within each b_max_ in order to demonstrate whether different methods make a significant difference given the same b_max_. Figure 3 shows the p-values between different b_max_ within each method, to see whether b_max_ made a significant difference in measures given the method. Additionally, Figure 4 shows p-values between DKI methods and DTI methods for MD and FA only, grouped by b_max_ of the DKI fits. We avoid the “multiple comparisons problem” by interpreting all results in the wider context, including the knowledge of how different fitting methods and b_max_ consistently affect measures across voxels within each subject.

**Figure 1:**
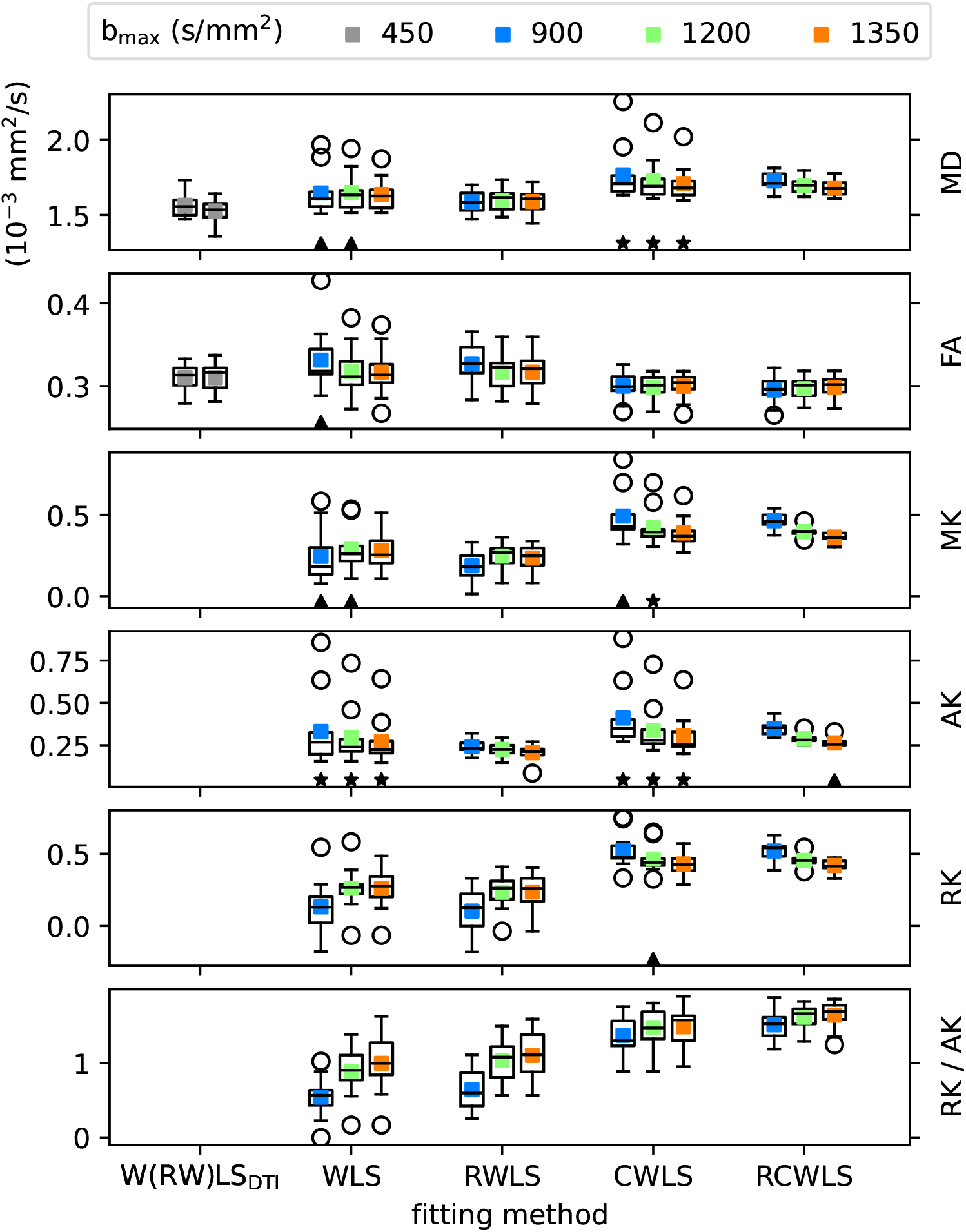
Average of DTI/DKI measures over myocardial voxels for all subjects. Colored points show the average over subjects, colored by b_max_. DTI fits W(RW)LS_DTI_ show WLS (left) and RWLS (right) together for convenience. The black triangles (stars) show where the Shapiro-Wilk test p-value ≤ 0.05 (≤ 0.01), i.e., the hypothesis of normality is rejected.

**Figure 2:**
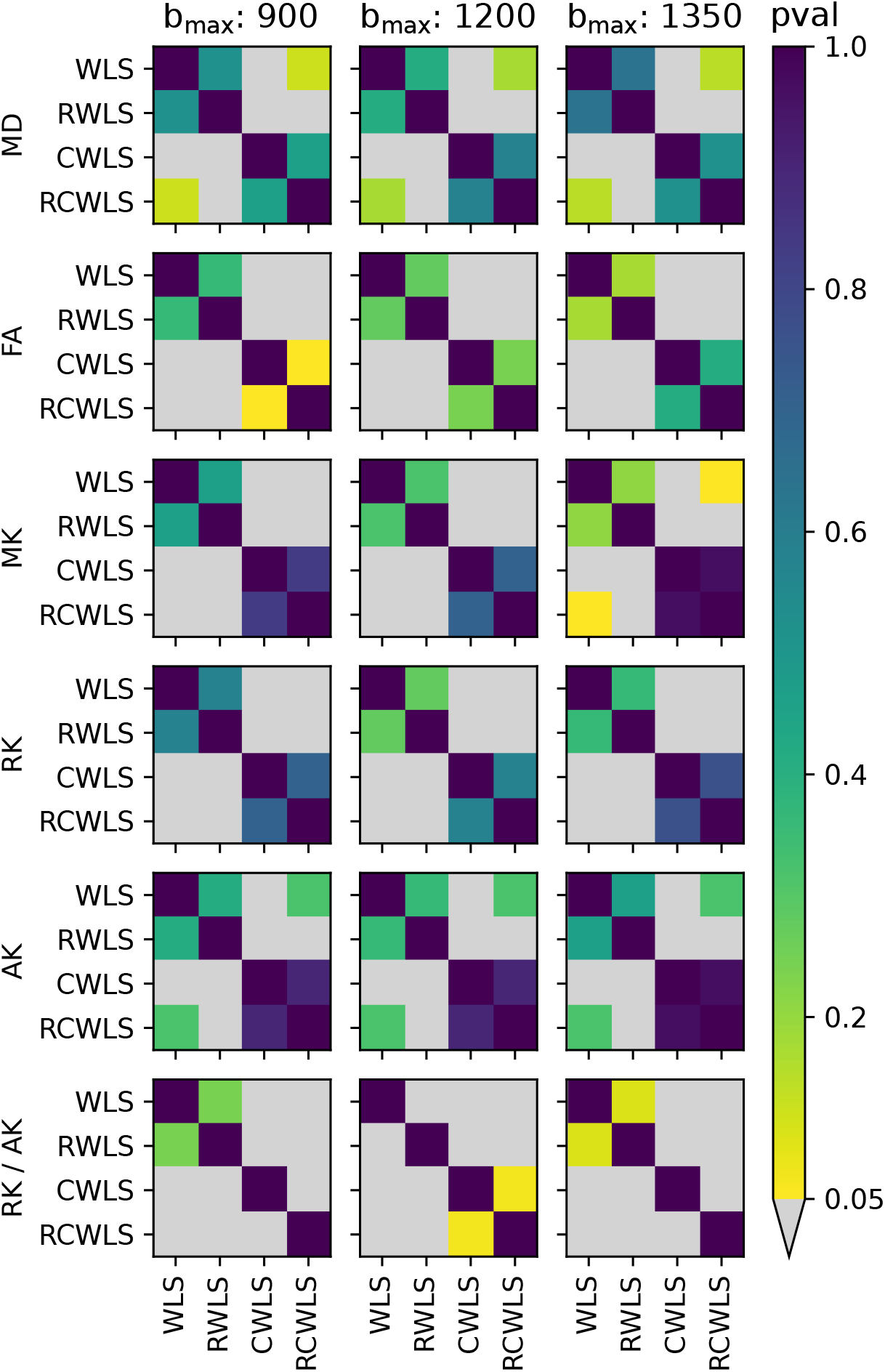
p-values from comparing DKI measures from different fitting methods given the same b_max_ (units s/mm^2^).

**Figure 3:**
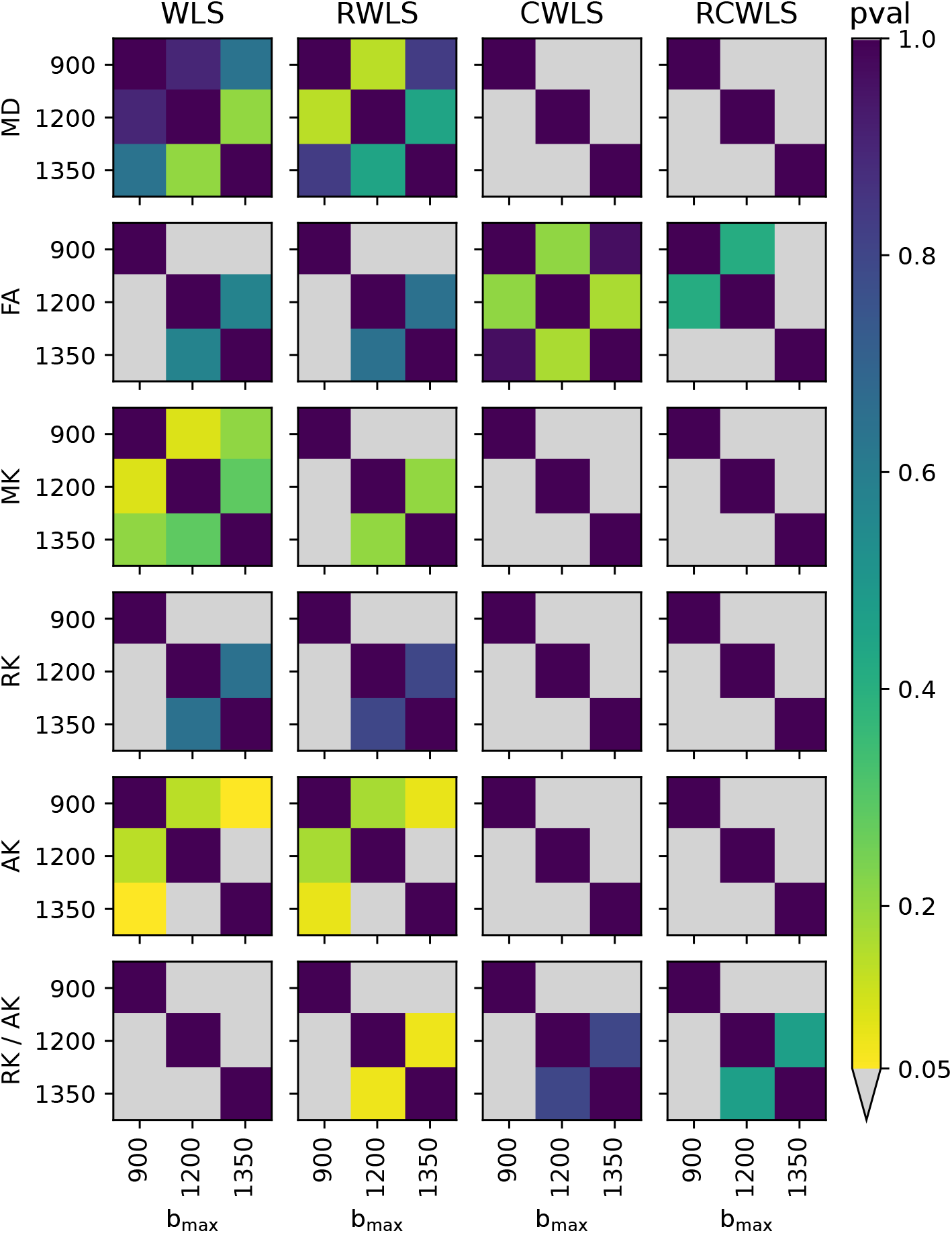
p-values from comparing DKI measures from different b_max_ (units s/mm^2^) given the same fitting methods.

**Figure 4:**
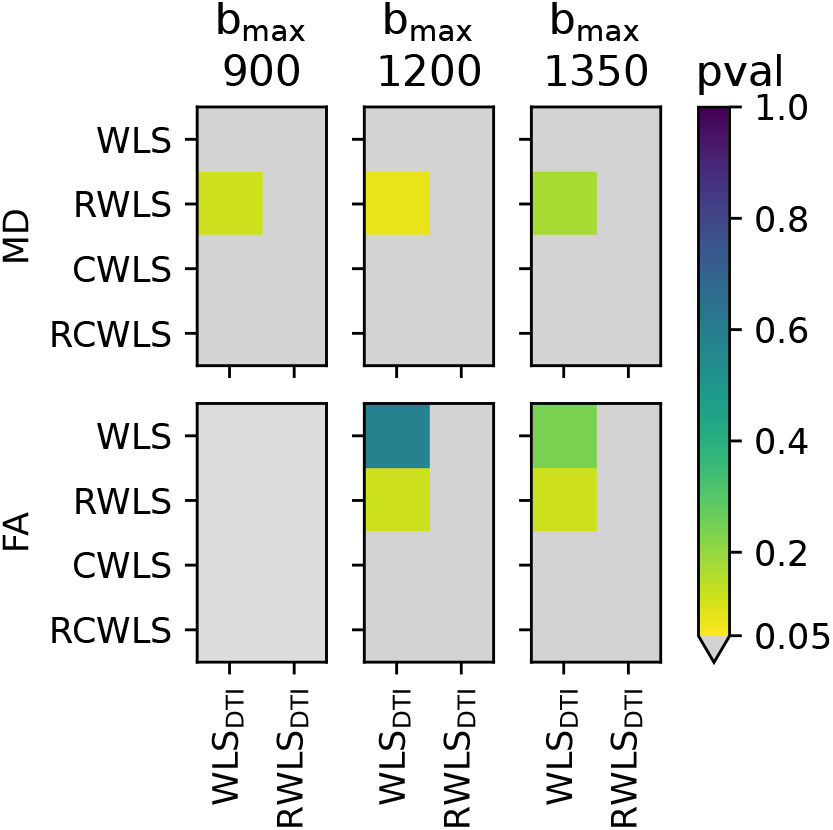
p-values from comparing MD and FA from DKI fitting methods for each b_max_ used for DKI fitting (units s/mm^2^) against DTI fitting methods (b_max_ = 450 s/mm^2^).

Adding constraints to a method, i.e., going from WLS to CWLS and from RWLS to RCWLS, made a statistically significant difference on all measures, for all b_max_ values, as shown in Figure 2. Figure 1 shows that constraints increased all kurtosis measures MK, AK, RK, and RK/AK, the latter results showing that constraints result in increasing RK more than AK. Increased kurtosis is consistent with the constraints ensuring that fitted parameters correspond to non-negative kurtosis. MK appears to increase with b_max_ for unconstrained fitting, but Figure 3 indicates this change is only significant going from b_max_ = 900 s/mm^2^ to b_max_ = 1200 s/mm^2^ and b_max_ = 1350 s/mm^2^, and only for RWLS but not WLS. For constrained fitting, MK, RK, and AK all decrease with b_max_, with Figure 3 showing these changes are significant for both CWLS and RCWLS. Nonetheless, Figure 1 indicates that the ratio RK/AK appears to increase with b_max_ for both constrained and unconstrained fitting. Note that the difference between RK/AK at b_max_ = 900 s/mm^2^ and b_max_ = 1350 s/mm^2^, and between RK/AK b_max_ = 900 s/mm^2^ and b_max_ = 1200 s/mm^2^, is statistically significant for both CWLS and RCWLS, while the difference between b_max_ = 1200 s/mm^2^ and b_max_ = 1350 s/mm^2^ is not (see Figure 3). However, given the significant trends with b_max_ for MK, AK, and RK, this is likely due to the subject with the outlier RK/AK value for RCWLS at b_max_ = 1350 s/mm^2^, as the inter-quartile range seems reasonably well separated.

In other words, the trend of increasing RK/AK with increasing b_max_ is likely to be meaningful.

MD increases from DTI (b_max_ ≤ 450 s/mm^2^) to unconstrained DKI (see Figure 4, which shows these differences are significant when comparing robust methods), and from unconstrained DKI to constrained DKI (see Figure 2, which shows these difference are significant except for comparing WLS, presumably due to the large spread of results for this non-robust method). For constrained methods, MD decreases slightly (but significantly; see Figure 3) as b_max_ increases. FA increases from DTI to unconstrained DKI (for robust DTI fitting, these differences are statistically significant compared with all DKI fitting methods; see Figure 4), but constrained DKI results in lower FA than DTI fitting (statistically significant for all DTI vs DKI comparisons; see Figure 4). For constrained results, Figure 1 does not seem to indicate a change in FA with b_max_, although Figure 2 indicates that for RCWLS the difference between b_max_ = 900 s/mm^2^ and b_max_ = 1350 s/mm^2^, and between b_max_ = 900 s/mm^2^ and b_max_ = 1200 s/mm^2^, is statistically significant. However, this could just be due to the outlier FA value for RCWLS b_max_ = 900 s/mm^2^.

Robust fitting by itself, i.e., going from WLS to RWLS, and CWLS to RCWLS, gave large changes in measures for some subjects (in particular, reducing MD and MK) but did not significantly change the mean measure values over subjects, except for RK/AK where the difference between CWLS and RCWLS was statistically significant for b_max_ = 900 s/mm^2^ (*p* = 0.0322) and b_max_ = 1350 s/mm^2^ (*p* = 0.0244); for b_max_ = 1200 s/mm^2^, the p-value was *p* = 0.0674, which is reasonably close to the significance cut-off that, in the context of the other tests, we can infer that robustness does in general significantly alter RK/AK even when constraints are already present. Robust fitting, by having large changes for some subjects, generally reduces the spread of the results. The black triangles and stars in Figure 1 indicate that non-normality is only found for non-robust methods but not for robust methods, with the *single exception* of AK for RCWLS b_max_ = 1350 s/mm^2^, for which *p* = 0.0381, but given that 36 tests for non-normality were performed for measures from robust fitting, this is likely just a result of the multiple comparisons problem.

### 3.2 Example maps

Figures 5, 6, 7, and 8 show example measure maps for 4 different subjects. The colormaps for MK and RK/AK were chosen based on the boxplots for RCWLS in Figure 1, and in order to help emphasize whether values are above zero (MK) or above one (RK/AK).

**Figure 5:**
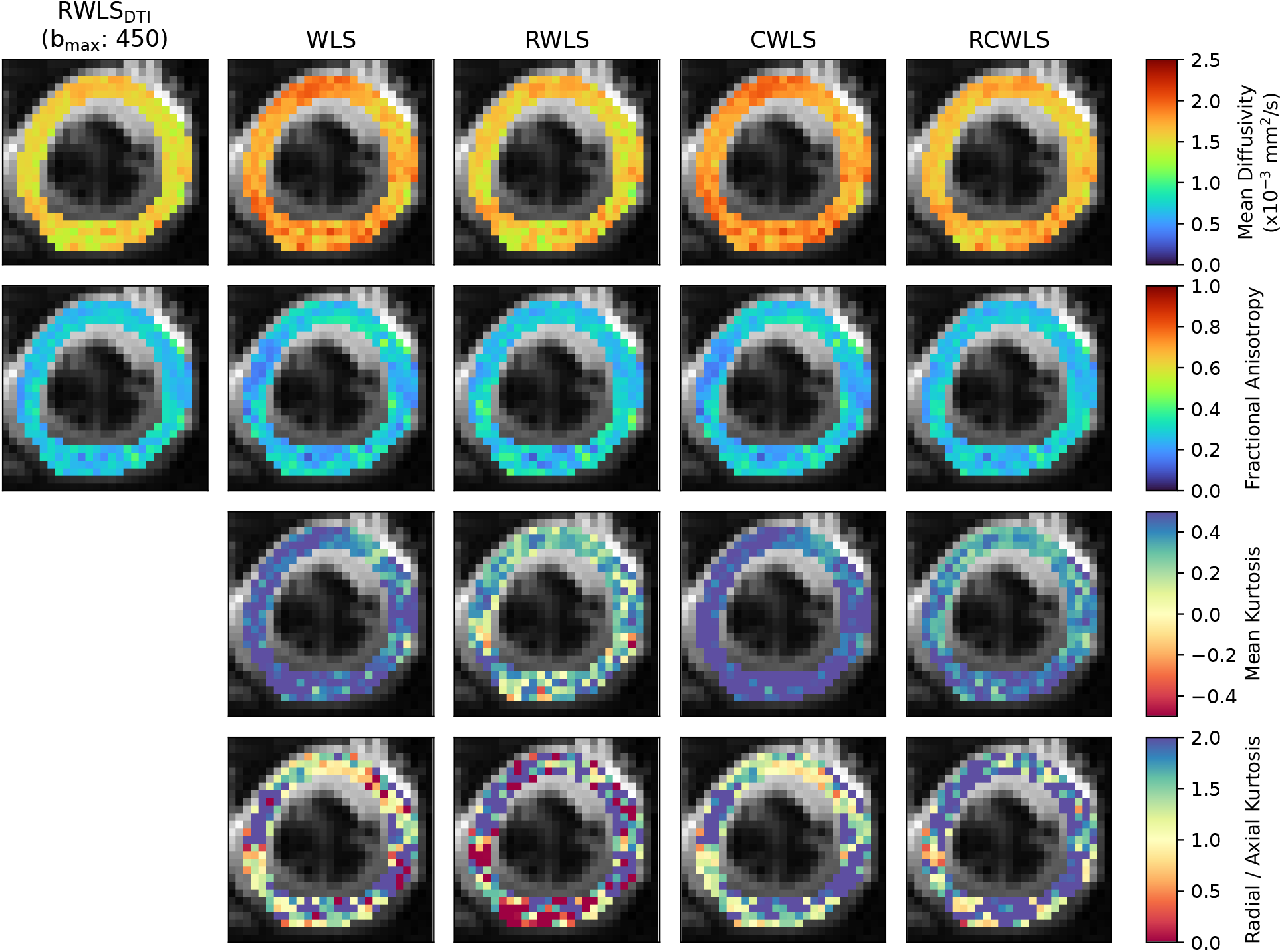
Maps for the mid slice of a single subject, for b_max_ = 1350 s/mm^2^. The first column shows MD and FA from DTI RWLS fitting (b_max_ = 450 s/mm^2^) for reference. All other columns show measures from DKI fits. In this example, robust fitting decreases MK, while constrained fitting increases MK. Using robust and constrained fitting (RCWLS) gives the most plausible results, with mostly RK/AK *>* 1 values.

**Figure 6:**
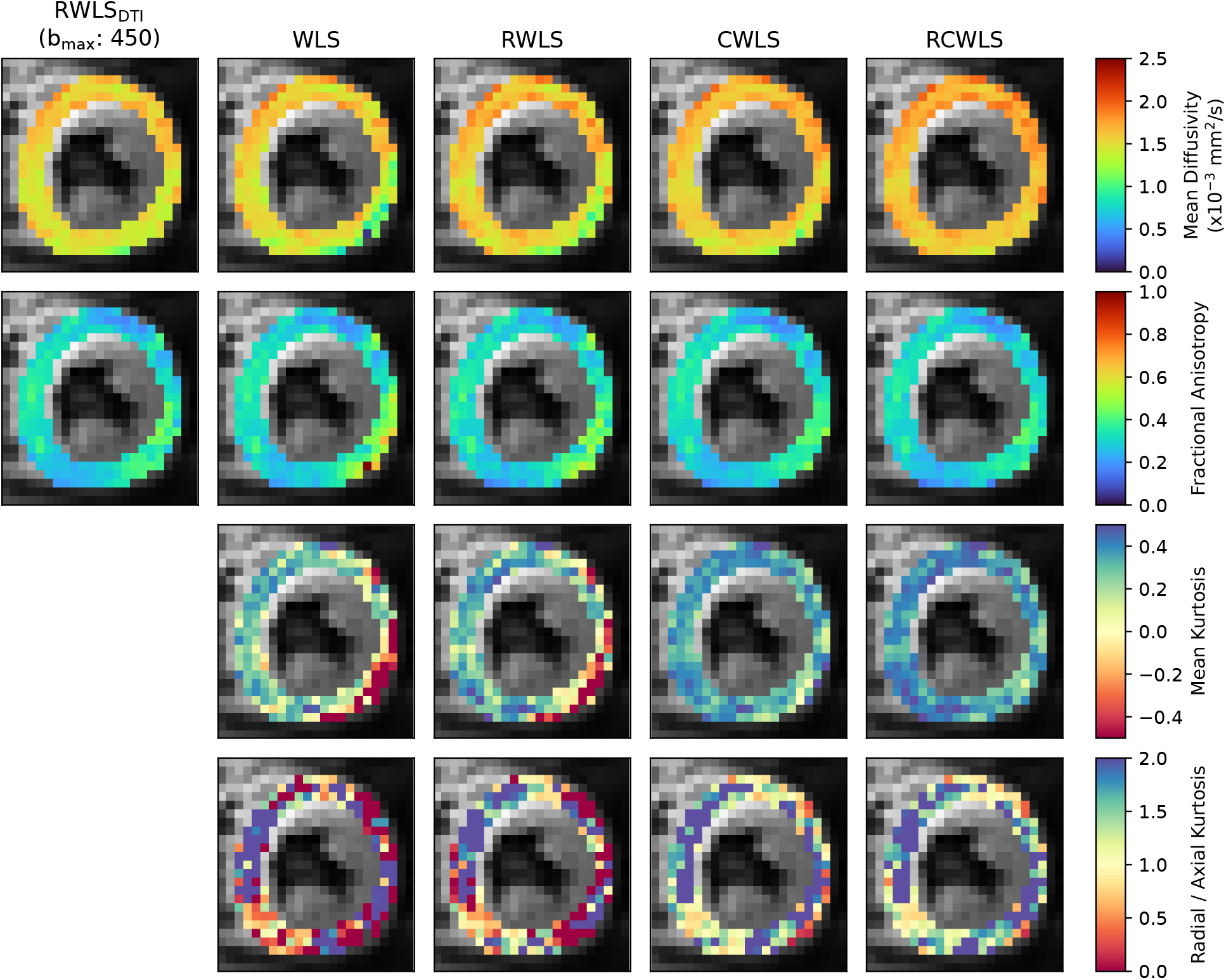
Measure maps for the mid slice of a single subject, for b_max_ = 1350 s/mm^2^. The first column shows MD and FA from DTI RWLS fitting (b_max_ = 450 s/mm^2^) for reference. All other columns show measures from DKI fits. Around 2 to 6 o’clock for the DKI fits is a region of implausibly low MD, high FA, and negative MK, for unconstrained fits. Constraints correct this region towards plausible values, with robust fitting visibly improving the region further in the same direction.

**Figure 7:**
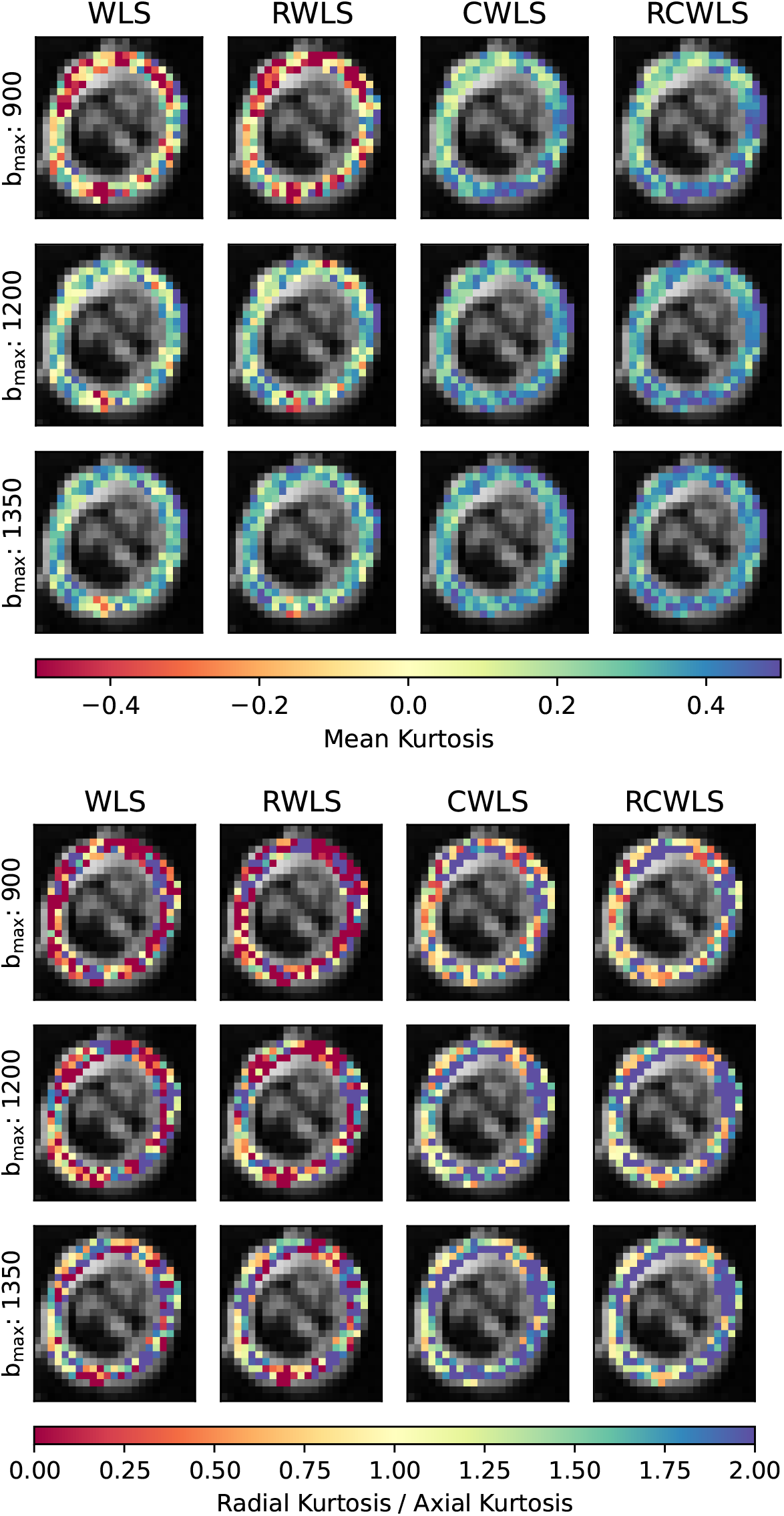
Mean Kurtosis (MK) and Radial Kurtosis / Axial Kurtosis (RK/AK) maps for the basal slice of a single subject, for all b_max_ values (units s/mm^2^) used for DKI fitting methods. Constraints increase MK given the same b_max_. For constrained fitting (CWLS and RCWLS), MK decreases with b_max_ in some regions and increases in others. RCWLS for b_max_ = 1350 s/mm^2^ has the most spatially homogeneous MK.

**Figure 8:**
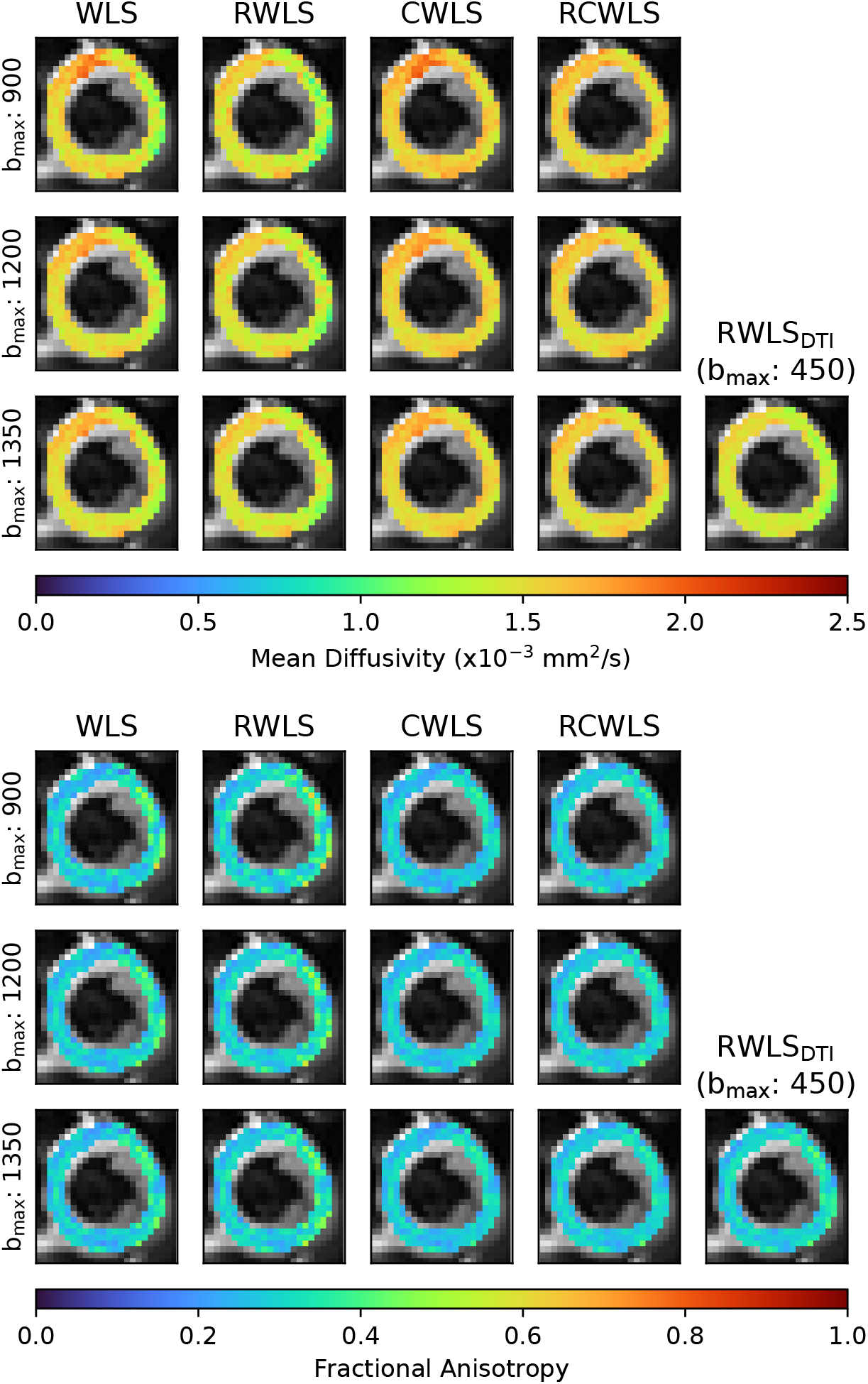
Mean Diffusivity (MD) and Fractional Anisotropy (FA) maps for the basal slice of a single subject, for all b_max_ values (units s/mm^2^) used for DKI fitting methods. Also shown are MD and FA from RWLS DTI fitting (b_max_ = 450 s/mm^2^) for reference. Robust fitting corrects elevated MD in the top-left (10 - 12 o’clock), but constraints are needed to correct the reduced MD and elevated FA on the right (12 - 5 o’clock).

Figure 5 shows basal-slice maps for b_max_ = 1350 s/mm^2^ for the subject with the second highest MD and MK for non-robust fitting WLS/CWLS in Figure 1. (The subject with the highest MD and MK values from non-robust methods shows a similar trend, but the images contain several artifacts that we excluded from the mean measures, making this subject a less suitable visual example). Robust fitting gave large changes over the whole myocardium, bringing measures into better agreement with other subjects, such that for robust fitting methods this subject is no longer an outlier in the boxplots of Figure 1. MD is reduced for robust methods vs non-robust methods, and FA is increased. Both MD and FA are more uniform for robust methods. Like for MD, MK is reduced by robust methods. Constraints increase MK and RK/AK, which is most noticeable between RWLS and RCWLS for the given colormap. In particular, RK/AK *>* 1 in nearly every voxel for RCWLS, and is relatively uniform. While WLS and CWLS also have RK/AK *>* 1 nearly everywhere, the values are highly heterogeneous compared to RCWLS. This example helps to make an important point: kurtosis can appear to be positive for non-constrained methods, but the kurtosis measures can still be corrupted. Going from WLS to RWLS results in a large reduction of MK, indicating that its consistently positive value for WLS was likely due to corruptions in the data (which, in this case, also caused an inflation of MD). By adding constraints to robust fitting, i.e. going from RWLS to RCWLS, consistently positive MK is recovered, but with much lower values than from the WLS fit.

Figure 6 shows mid-slice maps for b_max_ = 1350 s/mm^2^ where robust fitting had large effects in a localized region (around 2 to 6 o’clock). Most noticeable in this region are the low MD, high FA, and negative MK. In this region, both robust fitting and constraints independently increase MD, reduce FA, and increase MK, with RCWLS (i.e. both robust and constrained) doing so the most. The regions with RK/AK around 1 (yellow voxels) for RCWLS correspond with regions of relatively low FA (loosely interpreted as more similar levels of diffusion along the 3 primary axes of the diffusion tensor), which likely explains the reduced RK/AK in these regions. This example demonstrates that while robust fitting does correct some eroneous MD and FA values, constrained fitting makes a much bigger difference. Nonetheless, the effects of robustness in addition to constraints are still clear: RCWLS gives the most spatially uniform values.

Figure 7 shows MK and RK/AK basal-slice maps, for b_max_ = 900, 1200, 1350 s/mm^2^. Constraints give positive MK and make RK/AK *>* 1 overall. Differences between unconstrained and constrained fitting decrease with b_max_. Reading across the rows, constraints increase MK given the same b_max_. For RCWLS, MK decreases with b_max_ for most voxels, except a region between 8 and 12 o’clock which seems to have much lower MK values for b_max_ = 900 s/mm^2^. In this region, MK increases with b_max_. The result of increasing MK in some regions and decreasing in others as b_max_ increases, results in RCWLS for b_max_ = 1350 s/mm^2^ having the most homogeneous MK values across the myocardium. The pattern of RK/AK in this example follows the general trends in Figure 1, showing that both constraints and increasing b_max_ lead to increased RK/AK.

Figure 8 shows MD and FA maps in a basal slice, for b_max_ = 900, 1200, 1350 s/mm^2^, as well as MD and FA from RWLS DTI fitting (b_max_ = 450 s/mm^2^) for reference. Robust fitting corrects elevated MD in the top-left (10 - 12 o’clock), but constraints are needed to correct the reduced MD and elevated FA on the right (12 - 5 o’clock) - note that the region of reduced MD and elevated FA is also visible in the RWLS DTI results. Besides these regions, DKI fits slightly increase MD compared to DTI fits, and constraints also appear to generally increase MD further, which is in agreement with the overall trend across subjects. For FA, unconstrained DKI increases FA compared to the DTI FA, but constrained DKI reduces FA slightly compared to DTI, consistent with the overall trends. RCWLS results in the most uniform MD and FA, with the least variation with b_max_.

## 4 Discussion

### 4.1 The need for robustness and constraints

Our results show that both robust fitting and convexity constraints affect DTI/DKI measures in important ways.

Robust fitting can have large overall and regional benefits on individual subjects, which is invaluable for potential clinical applications. In the context of a study between different groups (e.g. healthy volunteers vs disease), the reduction in the spread of measures from robust fitting can lead to larger and more significant group differences [20]. Here, with a single exception likely from chance, only non-robust methods produced non-normal distributions of measures. Variation between subjects within a group ought to represent physiological variation (and non-gross measurement error) rather than the effects of image corruptions. Although the differences between non-robust and robust fitting were generally insignificant at the group level, the spread of the results of non-robust methods appeared to be responsible for these findings in several cases. Indeed, changes of MK with b_max_ were found to be significant for RWLS but not WLS, precisely because the MK values for several subjects were hugely inflated when fitting with WLS. For RK / AK there were significant differences between CWLS and RCWLS, indicating that changes from robust fitting on both RK and AK can result in a significant difference in their ratio. RK *>* AK is expected in myocardium, since there are fewer restrictions to diffusion parallel to the long axes of cardiomyocytes [3, 34]. Thus, our finding that robust fitting significantly increases this ratio is important.

Imposing convexity constraints resulted in statistically significant differences in all measures, for all b_max_ values. The precise changes, notably increased MK and RK/AK, are important. Only constrained fitting reliably shows RK *>* AK despite this being expected in myocardium [34] (for RCWLS b_max_ = 1350 s/mm^2^, RK/AK mean and median are 1.65 and 1.70 respectively). It is worth emphasizing that adding constraints does not just turn negative kurtosis into zero kurtosis in certain voxels (something that could be trivially obtained in post-processing the fitting results), but fundamentally alters the optimum coefficient vector (see examples Figures 5 - 6).

While it is known that MD and FA obtained from fitting the DKI signal representation can be different compared to the ones from the DTI signal representation [35], constraints further modify these values:

MD increases from DTI to DKI and from unconstrained to constrained fitting, but while FA increases from DTI to DKI, it decreases from unconstrained to constrained fitting, resulting in RCWLS giving the lowest FA values. This shows that the FA changes from DTI to DKI were largely in the context of parameter fits that violated constraints. To the best of our knowledge, other works noting differences in MD and FA between DTI and DKI have neither utilized constraints nor robust fitting.

RK/AK gives the most convincing evidence that using robustness and constraints together is important: the increase from CWLS to RCWLS is significant, as is the increase with b_max_ for RCWLS. Figure 5 helps show that the RCWLS results are not just the robust result with the negative kurtosis turned to zero - the kurtosis becomes convincingly positive, and all measures change when adding constraints. In this case, robust fitting reduced inflated MK caused by corruptions, while adding constraints increased MK. Figure 6 demonstrates a region where only robust fitting and constraints together appear to fully resolve a region of corrupted measures. In theory, RCWLS provides advantages greater than the sum of its parts: convexity constraints should make outlier identification easier, which should improve the weights in the final WLS fit.

### 4.2 b-value dependence

Varying b_max_ allowed for investigating the importance of constraints and robustness for different experimental designs: higher b_max_ data is more informative about kurtosis but has lower SNR. While previous work has noted the dependence of the performance of (simplified models of) cardiac DKI on b_max_ [3], it is noteworthy that MK and RK appear to increase with b_max_ for unconstrained fitting, but decrease with b_max_ for constrained fitting. This only serves to emphasize the importance of using fitting constraints. Due to these opposite trends, the differences between unconstrained and constrained fitting get smaller as b_max_ increases, suggesting fewer and milder constraint violations as data become more informative about kurtosis. In a few cases, significant differences between b_max_ = 900 s/mm^2^ and b_max_ 1200 s/mm^2^, and between b_max_ = 900 s/mm^2^ and b_max_ 1350 s/mm^2^, were found, while differences between b_max_ = 1200 s/mm^2^ and b_max_ 1350 s/mm^2^ were not. While this might suggest at first glance that the variation with respect to b_max_ is leveling out, note that the b_max_ values are not equally spaced.

We would like to point out that the results here are not time-normalized: higher b_max_ included all lower b-values, and so there is more data available for these fits. Our study sought to determine how the results changed as more information about kurtosis was available, specifically in the context of understanding the effects of robust fitting and constrained fitting, so the conflation from having both higher b-value data (which has lower SNR) and more data overall was not of particular concern. To perform an analysis on the appropriateness of b-values, we believe many more subjects would be required, as well as a more nuanced analysis of trends with b_max_ (rather than just paired tests). This work could be undertaken with the understanding that robust and constrained fitting ought to be used.

## 5 Conclusion

In this work, we have developed robust constrained weighted least squares (RCWLS), the first true robust estimation technique for DKI that incorporates necessary constraints on the signal behavior. Using in vivo human cardiac DKI data from healthy volunteers collected with a Connectom scanner, we determined that RCWLS is the most suitable fitting technique compared with others that lack either robustness or constraints.

## A Constraint matrices

Many basis elements can be excluded a priori because they cannot contribute, leaving:

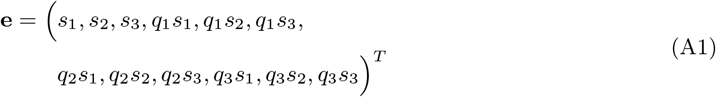

We can parameterize the matrix *H*(***θ***) as:

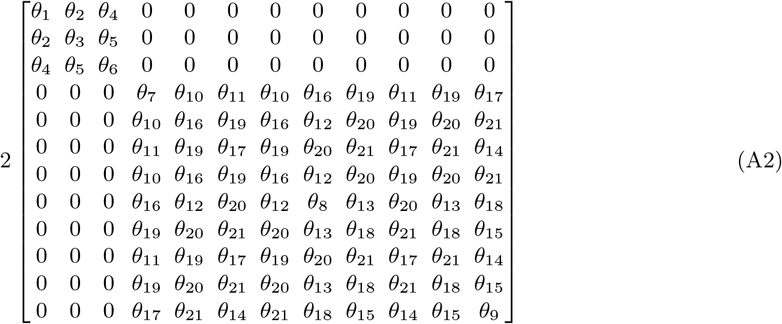

The ‘slack matrix’ *L*(***α***) is determined by Eq (7f). We can write 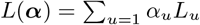, such that **e**^*T*^ *·L*_*u*_ *·***e** =0. There are 18 such matrices, so we parameterize *L*(***α***) as:

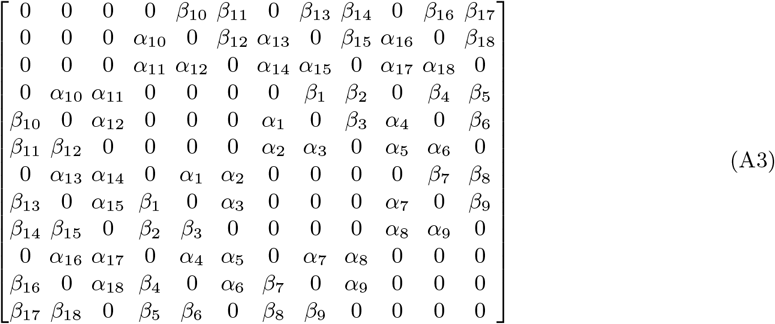

where *β*_*ij*_ = −*α*_*ij*_ (where *β*_*ij*_ are only used here to assist with display, they are not additional parameters).

## B Reduced constrained problem

As explained in the appendix of [18], the size of the semidefinite programming problem can be reduced (giving significant computational savings) by replacing Eq (7a) with

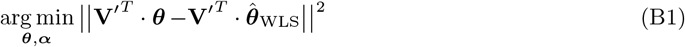

where 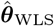 is the unconstrained WLS solution given by Eq (5) and **V** is given by the Cholesky factorization **X**^*′T*^ *·* **X**^*′*^ = **V** *·* **V**^*T*^. For RCWLS the problem is reduced at each iteration (the *unconstrained* fit result 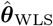 depends on the previous *constrained* fit). The semi-definite program Eq (7) only needs solving if the WLS estimate violates the constraints.

## Acknowledgments

S. Coveney would like to thank Serge Koudoro, Ariel Rokem, and Rafael Neto Henriques for help with implementing our work in DiPy, and Tom Dela Haije for advice on deriving the constraint matrices. This work was supported by Wellcome Trust Investigator Award grant 219536/Z/19/Z.

